# Dynamics and control of highly pathogenic H5 avian influenza in a threatened pelican population

**DOI:** 10.64898/2026.03.16.712014

**Authors:** Qiqi Yang, Olga Alexandrou, Ursula Höfle, Sara Minayo Martín, Serafeim C. Chaintoutis, Evangelia Moutou, Chrysostomos I. Dovas, Louise Moncla, Bryan T. Grenfell, Giorgos Catsadorakis

**Author notes:** These authors jointly supervised this work (Bryan T. Grenfell, Giorgos Catsadorakis).

## Abstract

The ongoing epizootic of highly pathogenic avian influenza (HPAI) continues to cause massive deaths in wildlife. Fundamental understanding of its disease ecology in natural populations is urgently needed. This knowledge has been hindered by the difficulty of acquiring data on epidemic dynamics. Here, using data collected from a threatened population of Dalmatian pelicans (*Pelecanus crispus*), we recover the epidemiological and evolutionary history of one of the largest HPAI wildlife mortality events. The results show that this devastating outbreak was likely seeded by a single introduction associated with movement of the species. By estimating epidemiological features of two consecutive outbreaks in the same population, we show that panzootic H5N1 since 2022 likely exhibits higher transmissibility and longer shedding time in non-reservoir birds, compared to previous H5NX subtypes. We also evaluate effectiveness of past and future control measures: carcass removal during the outbreak is shown to have surprisingly little impact on mitigating the mortality; and current H5 vaccines relying on capture and injection to deliver cannot establish herd immunity in a wildlife population. The results provide the first field evidence supporting the hypothesis that viral fitness difference of H5N1 to previous H5NX subtypes is the key cause of the expanded epizootic and panzootic since 2022, and on highly debated HPAI management strategies in wildlife populations.

**Author Summary:** Since late 2021, a panzootic of H5N1 highly pathogenic avian influenza (HPAI) has caused unprecedented mass mortality in wildlife. Many severely affected species are critical for ecosystem functions, including several threatened and endangered species. However, fundamental knowledge of HPAI disease ecology in natural populations is still lacking, and the effectiveness of potential controls is under debate. Here, using data collected from one of the largest HPAI outbreak in wild animals – over 1700 deaths (80% of the population) in a threatened population of Dalmatian pelicans in Greece, for the first time we recover the transmission dynamics of H5N1 in a migratory bird population. Based on the recovered dynamics, we show that removing carcasses during the outbreak was surprisingly ineffective, and future potential vaccination would require a novel delivery method to establish population immunity in wildlife. Our study provides new insight in the epidemiology of HPAI clade 2.3.4.4b in wildlife, and provides a foundation for assessing interventions within this complicated system.

## Introduction

Since late 2021, highly pathogenic avian influenza (HPAI) has caused unprecedented mass mortality in diverse avian and mammalian species globally^(1,2,3)^. This ‘panzootic’ has affected many critical species in ecosystems and several threatened and endangered species, impacting biodiversity and ecosystem health.

Regional and global surveillance and information sharing networks have made it possible to understand HPAI epidemiology, evolution and spread at large geographical scales. However, we have relatively little understanding of HPAI epidemic dynamics in individual wild bird populations. This gap is largely a question of data: time series of incidence or mortality data are required to quantify epidemics, but such detailed data are rare in imperfectly observed wild bird populations. To date, only one study has documented time series data of deaths during two HPAI outbreaks (H5N8 in 2016 and H5N6 in 2017), allowing the authors to infer epidemiological parameters in a wild mute swan population in the United Kingdom^(4)^. The authors fitted an epidemiological model with generation interval estimates from experiment studies. They inferred a basic reproduction ratio of infection, *R*_0_, of 2.25 [1.92*−*2.68] and 2.69 [1.40*−*5.5] from two H5 HPAI outbreaks in 2016 and 2017, respectively^(4)^. In addition, HPAI epidemics in individual domestic poultry flocks have been analyzed using population-level models, assuming frequency-based transmission^(5)^. Although there are no other models of HPAI in a single natural population, a few models were developed for low pathogenic avian influenza (LPAI) or avian influenza prevalence dynamics in a bird community of multiple species^(6,7,8)^. An epidemiological model was applied for LPAI prevalence dynamics within a season (from September 2005 to July 2006) in a local bird community in France; it incorporates environmental transmission through water and reflects a mix of frequency and density based transmission^(7)^.

The importance of host demography in shaping epidemic dynamics is well established in wildlife disease ecology^(9,10,11)^. For LPAI, models and empirical data suggest that annual births^(12)^ and migration^(8,13)^ are important factors in avian influenza prevalence and viral diversity^(14)^. For example, LPAI prevalence and seroprevalence data in multiple avian species in Australia suggest that seasonal bird frequency is associated with LPAI prevalence of the species. An epidemiological model was applied to LPAI prevalence dynamics in a migratory waterfowl community across post-breeding and migration sites in North America. It revealed the birds are likely infected with LPAI early in the fall migration^(8)^. For HPAI, phylodynamic models show that bird migration is highly relevant to HPAI spread and evolution at regional and global geographical scales^(15,16,17)^. Despite the importance of incorporating the seasonal population dynamics of avian hosts, these variables have not been included in HPAI modelling in an individual wild bird population. As HPAI becomes endemic in more wild bird species, understanding epidemics in individual populations with the incorporation of host demography will be key to design surveillance and to evaluate control measures.

Here, we analyse HPAI outbreaks in a seasonally breeding population of Dalmatian pelicans (*Pelecanus crispus*) at Prespa, Greece. In a severe outbreak in 2022 following a mild outbreak in 2021, 1734 individuals (over 40% of the south-east European population, or about 10% of the global population) was lost^(18)^, reversing conservation efforts in the past two decades^(19)^. In 2022, other colonies of Dalmatian pelicans in the region also confirmed HPAI-related mortality with a total of 128 deaths^(18)^, including Karavasta Lagoon in Albania, Skadar Lake in Montenegro^(20)^, and three of the Romanian colonies in the Danube Delta. Thanks to the long-term monitoring of the population for conservation purposes, demographic data and death counts were collected during the two outbreaks. This presents a rare opportunity to understand the epidemiology of HPAI in wild birds. Here, we begin by inferring the evolutionary origin and introduction time of the viruses using viral genome data and phylogenetic methods. With an epidemiological model, we then quantify the epidemiological parameters of HPAI H5 clade 2.3.4.4b in a wild bird population. Based on the parametrised model, we evaluate the effectiveness of current and potential control measures: carcass removal and vaccination^(21,22,23,24)^. Our study improves foundational understanding of avian influenza ecology in pelican species and provides insight on designing future control measures.

## Results

### Evolutionary source of the outbreaks

Geographical origin of the outbreak strains is suggested by genetically close sequences on the phylogenies (Fig 1 and Fig S7). In the 2022 outbreak, a genetically highly similar sample from a Dalmatian pelican in Albania (Karavasta Lagoon colony, Fig 1A) nests within the Greece outbreak cluster on phylogenies of all genes (Fig 1B and Fig S7). This implies the **direct transmission between infected pelicans in Albania and Greece**. However, due to the limited sequences, we cannot conclusively identify the dispersal direction between Lesser Prespa Lake and Karavasta Lagoon, nor at which colony (or a third site shared by the pelicans) transmission happened. Although traditionally the two populations were thought to not directly interact due to mountain barriers^(25)^, interactions are observed in our most recent tracking and ringing data (Fig S 4).

**Fig 1:**
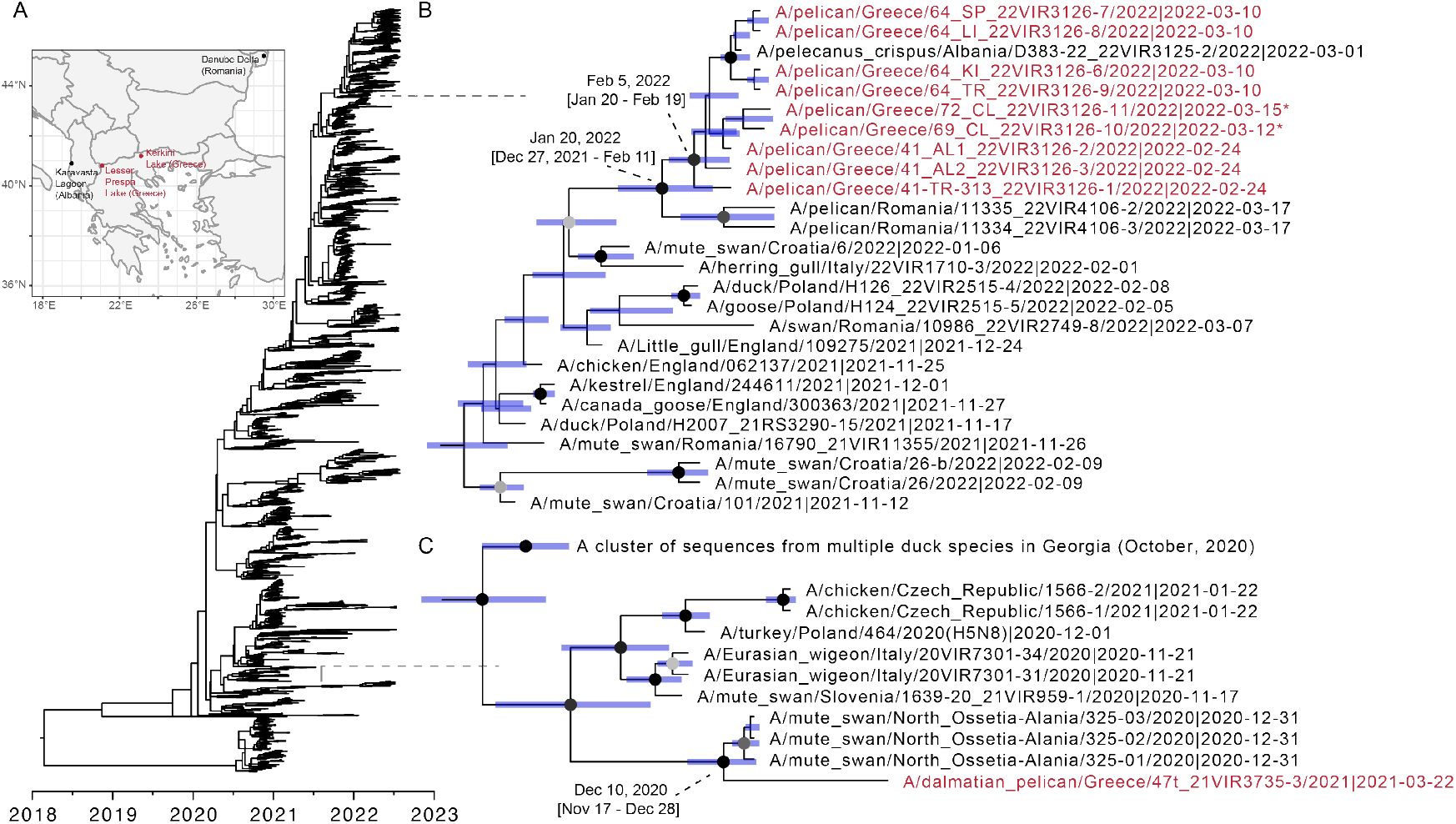
Bayesian time-scaled phylogeny of 4351 HA sequences. *(A)* Locations where samples from Dalmatian pelican are collected. *(B)* 2022 outbreak sequences and phylogenetically closest non-outbreak sequences. *(C)* 2021 outbreak sequences and phylogenetically closest non-outbreak sequences. The outbreak sequences are coloured in red. The divergence time of the outbreak cluster is labelled. The posterior time of the node is labelled by the blue bar. The nodes with posterior support greater than 50% are coloured from grey to black based on ascending support values. (*) marks the 2022 outbreak sequences that are sampled in other locations of Greece.

Two samples from Dalmatian pelicans in Romania (Danube Delta colony, Fig 1A) form a separate cluster and diverge from 2022/Greece outbreak samples on January 20, 2022 (95% HPD: December 27, 2021-February 11, 2022) (Fig 1B). The right tail of the divergence time largely overlaps with that of 2022/Greece samples. Besides, on phylogenies of other genes, the Romanian samples closely cluster with 2022/Greece samples (Fig S7). This body of evidence implies that the **HPAI infected pelicans in Romania and Greece likely acquired the initial infections from the same source**. Considering that direct movement between Prespa and Danube Delta population is extremely rare, their common wintering site, Kerkini Lake, could be where they acquired the initial infections. Additionally, a sample from a mute swan in Romania in November 2021 is closely related to 2022/Romanian and 2022/Greece samples on PB2, NP and NA phylogenies. This indicates that **the initial infection of Dalmatian pelicans was possibly acquired from other waterbird species in the same area**.

The 2021 outbreak strain clusters with three sequences sampled from mute swans in North Ossetia-Alania at the end of December 2020, with strong statistical support on the time-scaled HA phylogeny and ML trees of all other genes (Fig S7). The HA sequences of the mute swan sequences and Dalmatian pelican/Greece sequence diverge on December 10, 2020 (95% HPD: November 17-December 28), 20 days [3 days to 1.5 months] before the samples collected in North Ossetia-Alania (Fig 1C). This indicates **a direct link between HPAI-infected waterfowl in North Ossetia-Alania and the 2021 outbreak at Prespa**, and that the virus had been circulating in other waterbirds in the region since late winter of 2020 before infecting the Dalmatian pelicans at Prespa.

### H5 transmission dynamics

Integrating the transmission (Fig 2) and evolutionary dynamics (Fig 1), we reconstruct the history of the 2021 and 2022 epidemics (Fig 3), and summarise key epidemiology features.

**Fig 2:**
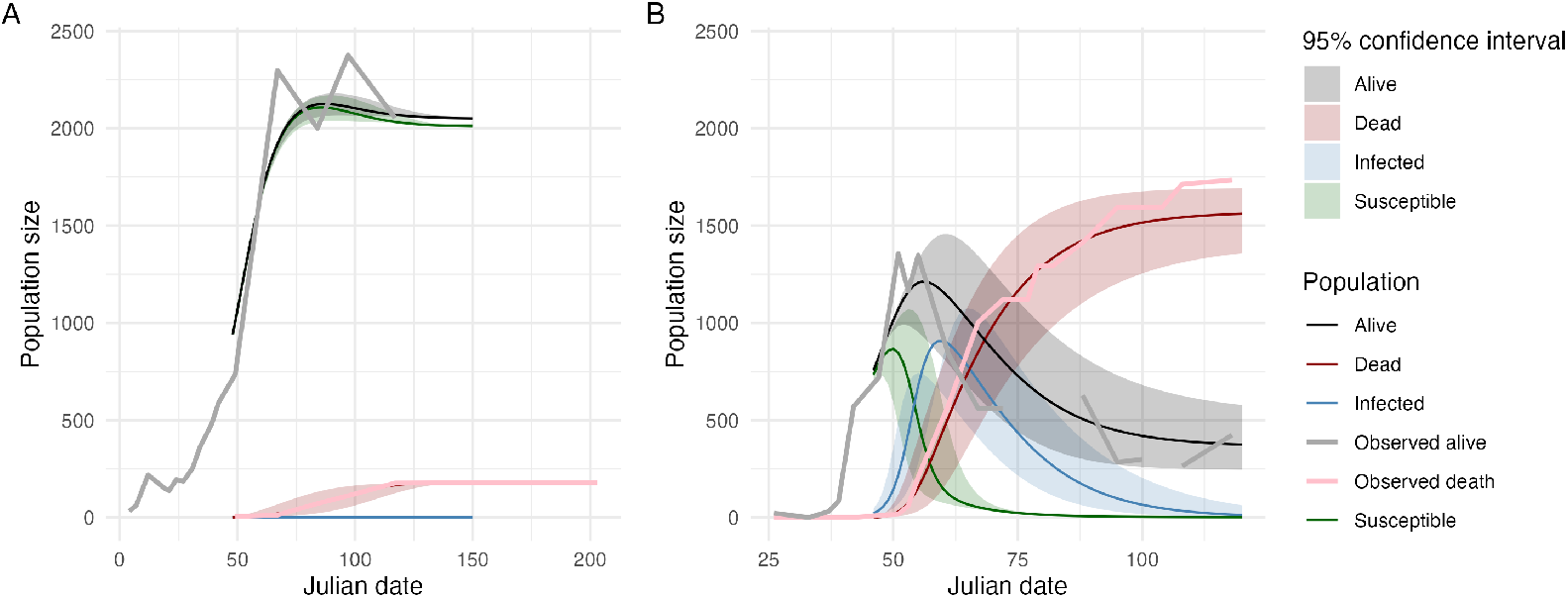
Predicted time series of the outbreaks, using parameters estimated by maximum likelihood method. *(A)* The 2021 outbreak. *(B)* The 2022 outbreak. Data of observed alive and dead birds are plotted using thick grey and pink curves, respectively.

**Fig 3:**
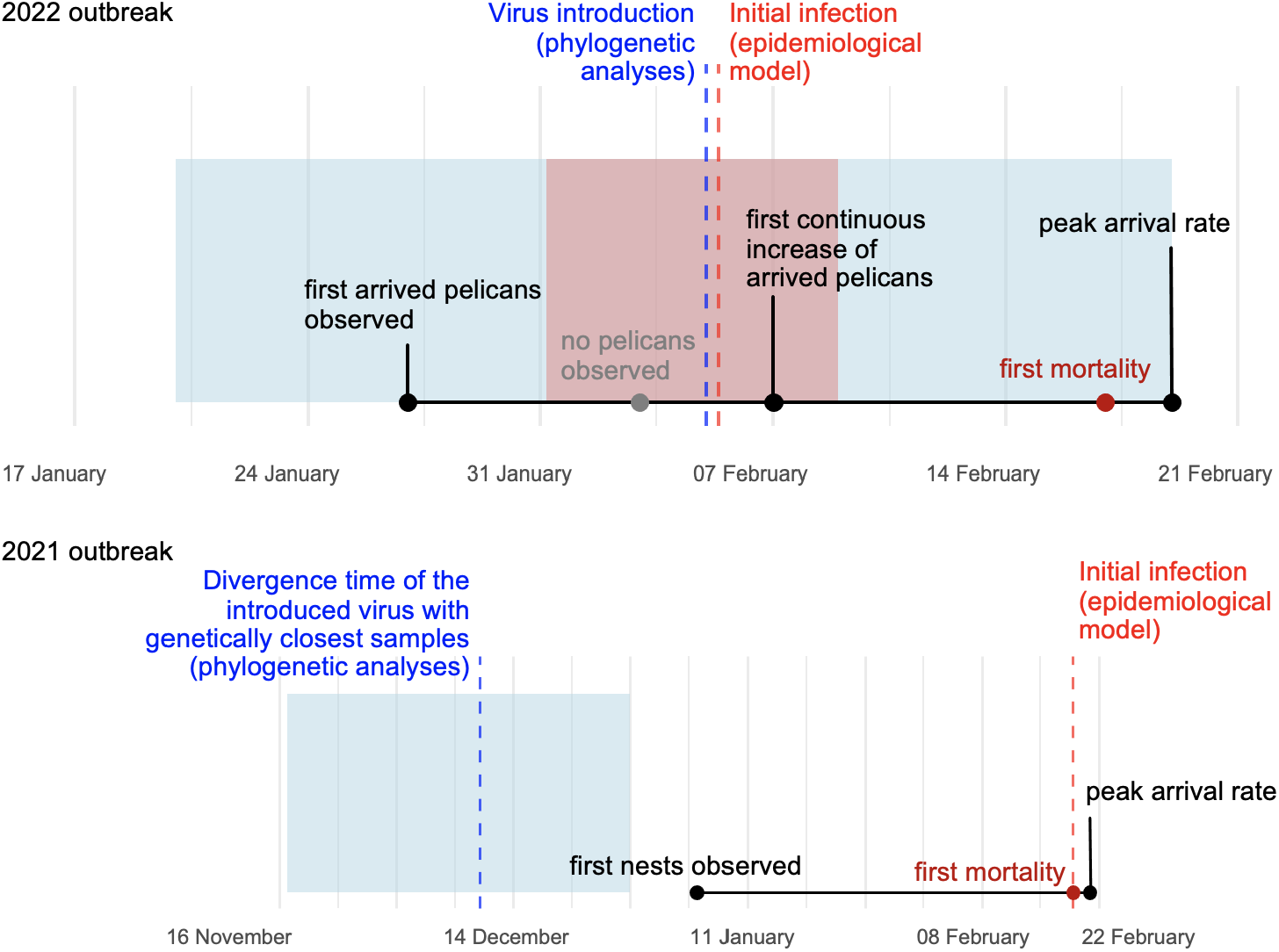
Epidemiology history of 2021 and 2022 outbreaks, reconstructed based on transmission and evolutionary dynamics modelling. The blue dashed lines indicate the inferred virus introduction time by phylogenetic analyses, the confidence interval of which is shown in a blue shaded box. The red dashed lines indicate the inferred initial infection time by epidemiological models, the confidence interval of which is shown in a red shaded box.

First, our fitted SIR model captures the overall outbreak progression pattern (Model 1). In particular, the observed dead and alive bird curves lie mostly within the model confidence limits for these quantities (Fig 2). The slight deviations are likely due to underestimated counts of alive birds during the early stage of 2022 and the rebound after massive nest abandonment in 2021.

The model-inferred epidemiological parameters (Table 1) indicate that the 2022 outbreak strain (EA-2021-AB) has higher transmissibility (*R*_0_ of 7 compared with 1) and longer infectious period than the 2021 outbreak genotype (EA-2020-A), while they have similar virulence (evaluated by fatality ratio). These estimates are consistent with the 10 times higher mortality of 2022 outbreak than 2021. Although virus challenge experiments comparing the two genotypes are lacking, previous experiments showed EA-2021-AB, when compared with EA-2020-C (another major genotype after EA-2020-A), causes more persistent viral shedding in Barnacle geese (*Branta leucopsis*) with more severe symptoms^(26)^.

**Table 1:** Epidemiological parameters inferred by maximum likelihood methods (details in Methods).

| Outbreak year | $R_0$ | Infectious period (day) | Fatality ratio | Initial infectious individuals/proportion |
| --- | --- | --- | --- | --- |
| 2021 | 1.01 [1.00-1.01] | 0.13 [0.11-0.16] | 82% [79%-85%] | 0.08 [0.03-0.3] / 0.009%[0.003%-0.03%] |
| 2022 | 7.07 [6.38-7.83] | 11.62 [8.04-16.81] | 81.2% [72.5%-87.6%] | 21 [10-45] / 2.8%[1.3%-5.9%] |

Second, the 2022 HPAI outbreaks in Dalmatian pelicans at Prespa and other locations in Greece were probably caused by a single introduction. This is reflected by the single statistically supported cluster of outbreak sequences on the time-scaled phylogeny of HA (Fig 1B) and maximum likelihood trees of other genes (Fig S7). Simulations of the epidemiological model also confirm a single introduction with 21 [10-45] or 2.8%[1.3%-5.9%] initial infectious individuals (Table 1), while repeated introductions cannot capture the observed epidemic dynamics. In the 2022 outbreak, the initial infected pelicans are among the early arrived ones. Phylogenetic analyses estimate that the single in-troduction occurred on February 5, 2022 (the median of the most recent common ancestor, TMRCA; 95% highest posterior density, HPD credible interval: January 20-February 19, 2022) (Fig 1B). This time perfectly overlaps with the infection time of the first death, estimated by the epidemiological model, February 5-6 (HPD credible interval: February 1-9, 2022, Fig 3). Unfortunately, similar inference of the virus introduction frequency and time cannot be concluded for the 2021 outbreak, due to the limited sequence data.

### Effects of control measures

#### Carcass removal

During the 2022 outbreak, to reduce transmission, conservation practitioners removed 1420 carcasses in total during March 17-24 and on April 11 and 15, 2022. To evaluate if carcass removal lowers the death toll in this case, we extend the simple SIR model to include infectious dead birds, which after the infectious period, become degraded (non-infectious) (Fig S 2B). We vary different ratio of transmissibility of dead birds versus the live infectious birds and infectious period. The best fit result estimates that the carcasses remain infectious for average 40 days and are similarly transmissible compared with live birds shedding viruses. Simulating the scenario of not removing carcasses with the parametrized model, we show that the **carcass removal does not affect total deaths of Dalmatian pelicans at Prespa colony** (Fig 4). Approximately 11 deaths (infected by carcasses) are averted; however, this is compensated by the increase of 11 deaths infected by alive birds.

**Fig 4:**
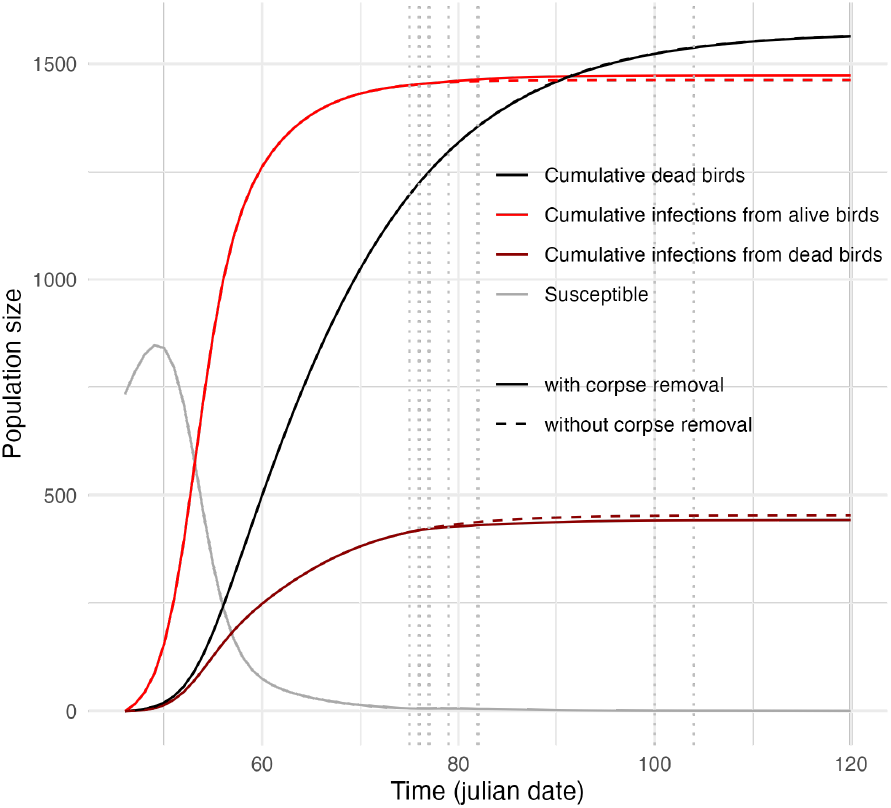
The comparison of simulated scenarios with (solid line) versus without carcass removal (dashed line). The model (Fig S 2B) is parametrized by best fits of the maximum likelihood method before carcass removal.

#### Vaccination

Avian influenza vaccination for natural bird population is typically considered impractical with current available vaccination strategies. However, emergency use of the vaccines could be considered for certain species, such as those of high conservation value and with high risk of being impacted^(21)^. For example, in 2023, emergency use of HPAI vaccine was approved for wild California condors (*Gymnogyps californianus*) in the US^(27)^. Recently, a HPAI H5 vaccine was shown safe and successful in inducing antibodies for captive Dalmatian pelicans and other 22 zoo bird species^(24)^. Transmission studies are still needed to investigate how the viral shedding reduction by HPAI H5 vaccines impacts transmission. Before experimental data are available, theoretical modelling can help evaluate the potential population-level outcome of vaccination^(28)^. Such evaluation is particularly important for vaccinating wild birds, as the optimal vaccination coverage, frequency and timing are all difficult and sometimes infeasible to achieve.

We simulate reactive vaccination in the parametrised model of the 2022 outbreak, assuming a ‘perfect’ vaccine that fully protects the individual from infection after one dose, the immunity of which lasts for the whole breeding season. For illustrative purposes, we assume that birds are vaccinated on a single day near the first infection being introduced. The results show that vaccinating 100% susceptibles any time after Day 52 would result in less than 100 deaths (Fig 5A), and any time before Day 72 returns more than 100 prevented deaths per 100 vaccinations (Fig 5C). Theoretically, the ideal vaccination time driving susceptibles below the epidemic threshold should be before the virus introduction, which would prevent the outbreak. However, only reactive vaccination is realistic in the context of injecting wild birds. For reactive vaccination, the most effective vaccination time to reduce HPAI impacts is Day 51-55 with between 147 and 151 prevented deaths per 100 vaccinated birds (Fig 5C). Interestingly, the ‘effectiveness’ peak time is only 4 days after the peak arrival time of birds in 2022. Our analyses suggest that if HPAI emerges during this time, mortality is the highest. This is consistent with the highest ‘effectiveness’ of vaccines if administered during this time. For Prespa colony, to avoid disturbing the breeding process, the only possible time to vaccinate the pelicans is before the peak arrival time (approximately Day 49 in 2022). Therefore, if vaccines were to be administered reactively to the full population, the best timing would be a few days before the peak arrival time. Realistically, it is infeasible to administer vaccine to approximately 10^3^ adult pelicans one by one. If vaccinating 50% adults (i.e. 500) on Day 50, the epidemic size decreases by approximately same size i.e. 500 ind. (Fig 5B and D). With a lower vaccination coverage, later timing of vaccination is favoured to decrease more deaths, despite the decrease in ‘effectiveness’ (Fig 5C).

**Fig 5:**
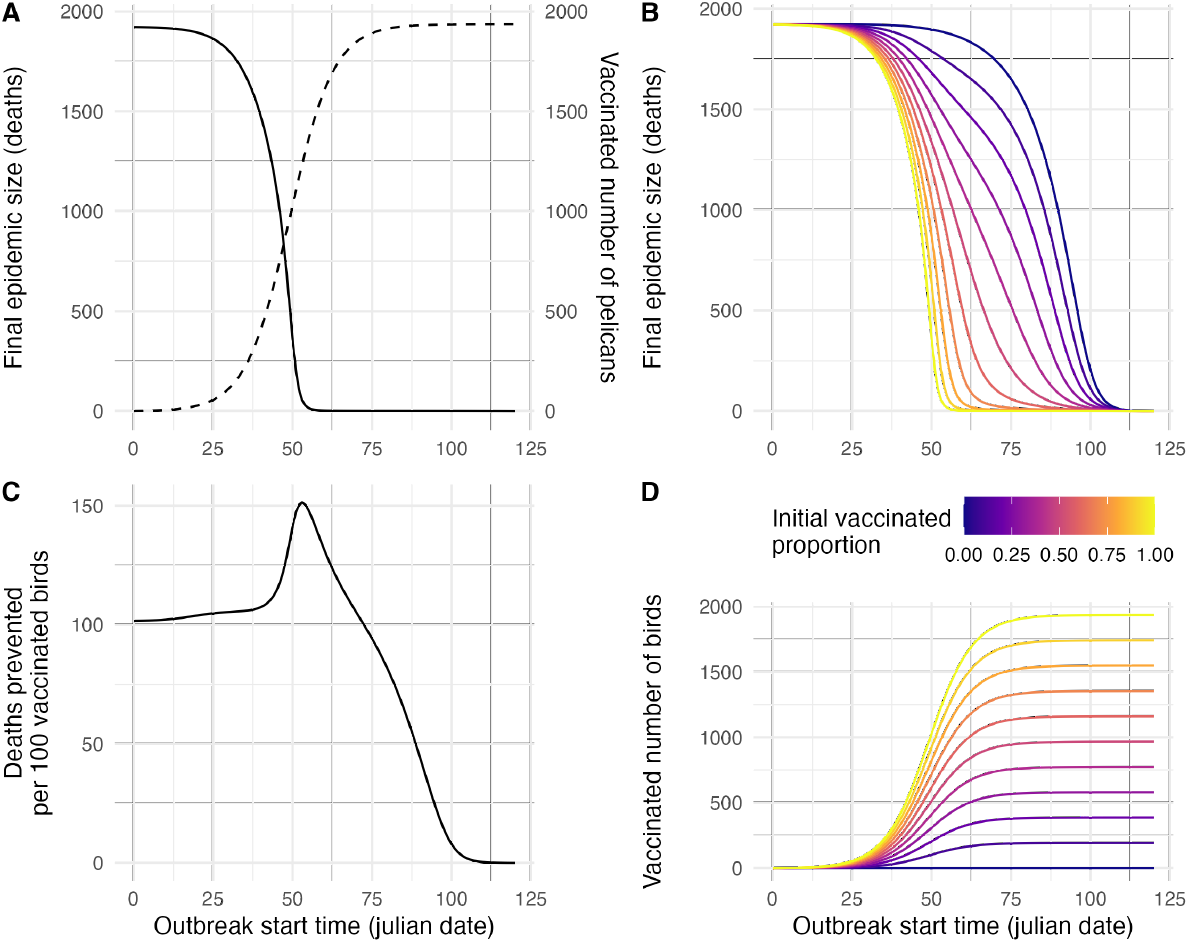
Simulated vaccination outcome. (*A*) The final epidemic size (solid line) and total vaccinated birds (dashed line), when vaccinating 100% susceptibles at different outbreak start time (x axis); (*B*) The final epidemic size when vaccinating a varying proportion (from 0 to 100% in colour) at different outbreak start time (x axis); (*C*) Deaths prevented per 100 vaccinated birds, when vaccinating 100% susceptibles at different outbreak start time (x axis); (*D*) The number of vaccinated birds, with a varying initial vaccinating proportion (from 0 to 100% in colour) at different outbreak start time (x axis).

## Discussion

Our epidemiological model successfully captures the important life history features of an HPAI outbreak in natural population, and demonstrates that for migratory birds, including bird seasonal immigration is vital for understanding HPAI dynamics and further evaluation of control measures. The viral phylogenetics reveal the potential geographical origin of outbreak strains, and highlight the same source of HPAI outbreaks in the European population of Dalmatian pelicans in 2022. More broadly, this phylodynamic combination of disease dynamics and viral phylogenetics helps build the foundation for the future epidemiological and phylodynamic modelling of HPAI in natural host populations.

### Causes of different outbreak outcomes

In 2021, death of 68 adults and 112 chicks was observed at Prespa population. Many of the chick deaths, however, could be due to loss of parental care during cold spell events. Therefore, HPAI may have caused up to 100 deaths. In contrast, the 2022 outbreak caused 1734 deaths in total (Fig S 5). The different outcomes of the outbreaks in 2021 and 2022 may also be affected by virus introduction time and population immunity, besides the outbreak strain difference in transmissibility (Table 1).

In 2022, the inferred virus introduction time and pelican arrival time largely overlap (Fig 3). In theory, this overlap could increase the virus transmission probability because during the initial stage of arrival, adult pelicans aggregate in a staging island and frequently interact (e.g., fight and flirt) with each other for mating. In contrast, the 2021 outbreak started after the mating time when the pelicans had dispersed and nested at all islets of the colony. Further research is needed to test this hypothesis.

Although 6 serological surveys were conducted in Dalmatian pelicans in Greece after the 2022 outbreak (Extended data table), we cannot conclude whether population immunity is a factor in generating different outbreak outcomes in 2021 and 2022, as we do not have AIV seroprevalence data for this species prior to 2022. Nevertheless, the serosurvey results supported the inferred fatality ratio of the 2022 outbreak, showing 20-30% seroprevalence of H5N1 AIV specific neutralizing antibodies in adult/immature Dalmatian pelicans in February 2023 and AIV antibodies in 2025 (Extended data table). These results suggest that some level of population immunity may exist. However, due to very limited sample size, conclusions cannot be drawn. Long-term serology surveys in more individuals of Dalmatian pelicans are needed to further understanding on population immunity.

### Limitations and future work on evolutionary origin of the outbreaks

With the limited virus samples during the outbreak at Prespa colony and in the nearby area (within Greece and beyond), we cannot completely reveal the geographical origin of HPAI in Dalmatian pelicans at Prespa colony. Our analyses confirm that Dalmatian pelican colonies in the region are closely linked with the outbreak. Direct transmission between breeding populations in different colonies, as well as transmission at shared wintering sites of different breeding populations, are both possible routes as confirmed by evolutionary analyses. As there were no poultry outbreaks reported before the outbreaks in Dalmatian pelicans, HPAI was likely introduced from other waterbird species. However, due to the lack of virus samples in other waterbird species in the area during the outbreak time, we cannot confirm the exact infection sources. Improving avian influenza surveillance in waterbirds at major habitats in this historically undersampled region and establishing a regional surveillance network would be helpful for outbreak preparedness and furthering knowledge of virus dispersal.

We also observe that the outbreak time of both years are strikingly similar, despite the arrival timing of pelicans differs by one month and the ecology of the migration is very different between the two years due to weather events. This is consistent with a previous study of two consecutive HPAI outbreaks in the wild mute swan population in England^(4)^, supporting that the disease ecology of HPAI in wild birds may be more consistent than expected.

### Role of pelican species in HPAI ecology and evolution

Different species of pelicans, including great white pelicans (*Pelecanus onocrotalus*), and Peruvian pelicans (*Pelecanus thagus*) have been severely affected by HPAI infection in Africa^(29,30)^ (Outbreak #1 on Fig S 6, Central Eurasia^(31,32)^ (Outbreak #2 on Fig S 6, and South America^(33)^. Besides, previous research shows pelicans have high risk of being exposed to HPAI^(15)^. Despite impacts of HPAI on pelican species, avian influenza epidemiology in wild pelican populations is unclear, largely due to the difficulty of capturing pelicans and conducting sampling.

The recent outbreaks caused by clade 2.3.4.4 is not the first time Dalmatian pelicans have been affected by HPAI. In 2014/15, clade 2.3.2.1, the only clade prior to clade 2.3.4.4 that has caused widespread mortality^(15)^, caused over 100 deaths of Dalmatian pelicans at Danube Delta in Romania and 20 deaths in Bulgaria^(32,34)^. Interestingly, the 2022 outbreak variant originates from HPAI viruses sampled from Dalmatian pelicans in September 2021 in south western Russia close to Kazakhstan (Saratov Oblast) (Fig S 6). The 2021 outbreak genotype also was first detected in the same area (Chelyabinsk Oblast) in July/August 2020. In the same region (Chelyabinsk, Kurgan and Tyu-men oblasts), an outbreak occurred from May to July causing 50% mortality of Dalmatian pelicans (decreased from 1200-1400 pairs to 600 pairs)^(31)^ (Outbreak #2 on Fig S 6). Whether the variant went through positive selection and became more adapted to Dalmatian pelicans is an important open question.

It remains unclear why great white pelicans that breed side by side with Dalmatian pelicans were not affected by HPAI outbreak in 2022. Our serological survey of great white pelican nestlings after the 2022 outbreak show that they are all seronegative, suggesting that they were not exposed to avian influenza virus during the outbreak. As the first mass arrival of great white pelicans is in early April at the later stage of the outbreak, it is likely that the virus wasn’t successfully spreading in the population. It is also possible that the long-distance migrant adults have immunity to HPAI H5 clade 2.3.4.4b due to previous exposure in the winter of 2020 - as reported from Senegal and Mauritania in Africa, a spillover from nearby chicken farm caused thousands of deaths of great white pelicans, with approximately 8% of the population in Senegal. The interesting pattern is that juveniles die at a distinctly higher ratio than adults: 740 juveniles and 10 adults in Senegal, 2081 juveniles and 59 adults in Mauritania^(29,30)^. Therefore, it is likely that adult great white pelicans have acquired some level of immunity from previous exposure. The pre-existing immunity may explain that great white pelicans never were affected in three different years, even though a few (4-12 ind.) great white pelicans always nest side by side to the early arrived Dalmatian pelicans. It is also possible that these individuals of great white pelicans have active infections and transmit the virus to Dalmatian pelicans. These hypotheses would need further virus detection and serology surveys in great white pelicans.

### Implications for evaluation of control measures

Currently, there are no standardised measures for controlling an active avian influenza outbreak in wildlife population^(21)^. The effectiveness of potential control measures, for example, removing carcasses, is under debate and discussion. During the 2022 outbreak at Prespa colony, a huge amount of human and physical resources were used to remove 1420 (82% of the total) carcasses. We show that in this specific case, the overall impact of removing carcasses in reducing deaths or infections is likely to be small. Although this measure lowers the deaths infected by carcasses, it is compensated by the deaths infected by alive birds. Although carcass removal was ineffective in controlling the 2022 outbreak in the Dalmatian pelican population at Prespa colony, it could be effective for other outbreaks - the effect is highly associated with outbreak epidemiology, and should be analysed case by case. In addition, carcass removal likely has other important positive impacts, for instance, reducing the contamination of the environment and preventing or reducing infections of other species. Scavenging is one of the most important transmission routes of HPAI in wild animals, which is likely to be inhibited by carcass removal. Further, for habitats shared with humans, removing carcasses likely reduces risk of spillover to domestic animals and zoonotic transmission.

Vaccination is another potential control measure especially for highly vulnerable protected species. We predict the population-level outcome of a ‘perfect’ vaccine and the results show the complexity of vaccinating a wild bird population, e.g., the optimal timing may be related to population demography. Therefore, when the immunogenicity and the impacts on transmission of avian influenza vaccines are tested, the population-level outcome of vaccine should be evaluated using theoretical models that are validated with long-term demography data and the immune landscape data of the population. Our results also highlight the challenges of vaccinating a wild bird population. Currently, HPAI vaccines are administered by injection to wild or captive birds. To inject a wild bird, the first step is to capture it. Given the extreme difficulty of capturing pelicans in the wild, it is impossible to achieve any effective vaccination coverage as predicted by our model even when the vaccine is ‘perfect’ (Fig 5). Furthermore, in reality, HPAI vaccines do not provide ‘perfect’ immunity; two doses of the vaccines are usually needed to provide strong and durable immunity^(23,24,22)^. Administrating the same wild pelican with two doses of vaccine means capturing the same individual twice during a few weeks or months, which is impossible to achieve. Therefore, the only possibly effective HPAI vaccines for a natural wild bird population especially large water birds like pelicans, would need novel administration routes that do not require capturing birds and can be scaled up to the population level, such as oral rabies vaccines for wildlife^(35)^ and a recently developed aerosol avian influenza vaccine^(36)^.

In conclusion, our study advances the understanding on the ecology and control of highly pathogenic avian influenza epidemics in a natural population. More broadly, it underlines the importance of integrated evolutionary epidemiological and immunological surveillance for understanding ‘panzootic’ (and hence zoonotic) threats across key wildlife species.

## Materials and Methods

### Population and epidemiological surveillance

#### Brief overview of the colony

Lesser (Mikri) Prespa Lake in North-Western Greece hosts the largest breeding colony of Dalmatian pelicans in the world^(19,37)^. These birds winter close to their breeding grounds and usually migrate to wetlands in north/northeastern Greece and western Turkey^(38,39)^. They start arriving at Prespa colony in January or early February and leave between mid-late June until October. During the breeding period, many of them forage for fish in wetlands outside Prespa, reaching even Kerkini Lake, 200 kms away (Study “Dalmatian pelican Greece (SPP data 2012-)”, ID: 9165543 on Movebank, movebank.org). Besides Kerkini Lake, other breeding colonies that have interaction with Lesser Prespa Lake include Cheimaditida Lake and Karla Reservoir. All of these colonies belong to the eastern European subpopulation and were all affected during the 2022 outbreak. In contrast, the two western colonies in Greece, which belong to the western European subpopulation, had no deaths of pelicans recorded^(18)^.

Besides Dalmatian pelicans, Lesser Prespa Lake hosts more than 25 species of migratory and resident water birds, including species of cormorants, geese, ducks and gulls^(40)^. A large number (approximately 721*−*779 pairs in 2020) of Great white pelicans breed side by side with Dalmatian pelicans on the colony^(41)^. They are long-distance migrants that fly through many important stopover sites on their southward migration to African wetlands along the Rift Valley to winter in eastern Africa^(38)^. The mass arrival of Great white pelicans is always later than Dalmatian pelicans, while their individual arrivals can be earlier.

#### Data collection during outbreaks

Using a mixed method of vantage point observation and drone photography, conservation practitioners and scientists at the Society for Protection of Prespa (SPP) have monitored and documented the demography of the Prespa population for the past decade. The breeding pairs are estimated to be 1585 in 2020 ^(42)^ and 991*−*1522 pairs between 2004 and 2010 ^(19,43)^. During the outbreaks in 2021 and 2022, the same methods are applied for counting the population size, nest number, and dead birds over the time. Because vantage point observation usually underestimates the counts at the colony, we use the ratio of the counts by vantage point observation versus the counts on drone images to adjust the vantage point collected data.

Using sterile swabs, oral and cloacal samples were collected from dead pelicans with the assistance of local veterinary authorities. Detection, diagnosis, and further sequencing of detected viral genome were conducted by the National Avian Influenza Reference Laboratory of Greece. The sequences were then submitted to Global Initiative on Sharing All Influenza Data (GISAID)^(44)^.

#### Active surveillance post outbreaks

Samples were collected from apparently healthy Dalmatian pelicans during ringing and radiotagging activities at major Greek pelican colonies, with leg-hold traps or hand catching using baits. From each bird, one cloacal and one oropharyngeal sample were collected, following protocols described previously^(45)^, using sterile swabs and stored in DNA/RNA shield (Zymo Research, CA, USA); and 1-2 mL of blood was collected from the toes and stored in heparinized sterile containers.

Oropharyngeal and cloacal swabs were screened in pools of five according to sampling date and location. For samples collected before December 2024, genetic material was extracted manually using the High Pure RNA isolation kit (Roche, Madrid, Spain). The extracted RNA was tested for AIV using matrix-gene real-time RT-qPCR as previously described^(46)^ and modified^(47)^ on a CFX96 real-time PCR detection system (Bio-Rad, Richmond, CA, USA). For samples collected since December 2024, RNA was extracted manually as previously described^(48)^, and was tested for AIV using matrix-gene real-time RT-qPCR as previously described^(49)^ on a CFX96 real-time detection system (Bio-Rad, Richmond, CA, USA). For PCR-positive samples, H5-specific PCR was conducted using primer sequences described previously^(50)^.

For serum samples collected before December 2024, they were heat inactivated for 30 minutes at 56 ºC and analyzed using a competitive ELISA (Ingenasa-Eurofins, Madrid, Spain) following the manufacturer’s instructions (Sensitivity, Se=0.97, Specificity, Sp=0.98). Competitive ELISA was also conducted for serum samples collected since December 2024 using IDvet NP ELISA kit (Montpellier, France; Se=0.97, Sp=1). Samples positive for antibodies against AIV in the competitive ELISA were subsequently subjected to a hemagglutination inhibition (HI) assay to detect the presence of antibodies against H5/H7 AIV serotypes (Se=0.85, Sp=0.99). The procedures were performed in accordance with the Manual of Diagnostic Tests and Vaccines for Terrestrial Animals (OIE, 2023). For serum samples before December 2024, reference antigens H5N2 (A/Ost/Den/72420/96) and H7N7 (A/Tky/Eng/647/77) reference antigens from the Animal and Plant Health Agency (APHA, Weybridge, United Kingdom) laboratory were used; and HI positive samples were retested for confirmation using reference antigen H5N1 (A/Chick/Scot/19/59). For serum samples since December 2024, reference antigen H5N1 (A/mallard/Italy/3401/05) was used.

#### Ethical considerations

Official permissions for capturing and sampling of pelicans were provided by the Hellenic Ministry of Environment and Energy (74506/2354/20-7-2022, 123240/4001/6-12-2022, and 9270/324/28-1-2024). All activities related to capture, handling, and sample collection in wild pelicans were carried out by experienced personnel and according to European and national legislation.

### Viral genome data and phylogenetic analyses

#### Sequence data

Full genomic sequences generated from 10 samples from dead pelicans in the 2021 and 2022 outbreaks were all HPAI clade 2.3.4.4b and differed in subtypes: H5N8 for the 2021 outbreak and H5N1 for the 2022 outbreak. It should be noted that only one sample was sequenced from the 2021 outbreak, and seven of nine samples from 2022 were from the breeding population of Dalmatian pelicans at Prespa, Greece while the other two are sampled from Dalmatian pelicans in two other locations (Kerkini Lake and Aggelochori, Thessaloniki) in Greece.

For background sequences, 4351 complete HA sequences of H5 Influenza A viruses sampled from avian hosts in Africa, Asia and Europe (country-level counts are summarised in Fig S 1) between July 1, 2020 and August 1, 2022 with complete collection date, were downloaded from GISAID in June 2025. The other gene segments of the 4351 viruses were also downloaded. The alignment of sequences were conducted using MAFFT^(51)^.

#### Maximum likelihood trees

To select an optimal model for nucleotide substitution, standard model selection method (similar as jModelTest^(52)^) was conducted using IQTree v3 ^(53)^. For the HA dataset, the optimal model is selected to be a general time reversible (GTR) model^(54)^, with gamma distributed rate variation among sites^(55)^, a proportion of invariant sites, and the empirical base frequency. Optimal nucleotide substitution models were selected for other gene segment datasets using the same method. The maximum likelihood trees were constructed using IQTree v3 ^(53)^. The standard nonparametric bootstrap was used for assessing branch support.

#### Bayesian time-scaled phylogeny

The strict clock model was selected as the molecular clock model for time-scaled phylogeny inference, given its robustness to sequence sub-sampling or selection^(56)^. The time-scaled phylogeny, the molecular clock model, the nucleotide substitution model (GTR + F + G4 + I, as described in ‘Maximum likelihood trees’) and the coalescent model (constant population size) were jointly estimated using a Markov chain Monte Carlo (MCMC) approach implemented in BEAST v1.10.5 ^(57)^ with BEAGLE^(58,59)^. The MCMC chain was run for 400 million generations with burin-in of 10%, sampling every 40000 steps. Convergence of the MCMC chain was checked with Tracer v1.7 ^(60)^. From the posterior distribution, we sampled a maximum clade credibility tree, and the time of most recent common ancestor of sequence clusters.

### Epidemiological model and data fitting by maximum likelihood estimation

#### A simple epidemiological model

To estimate epidemiological parameters of the 2021 and 2022 outbreak, we modified a classic Susceptible-Infectious-Recovered (SIR) model^(61,62)^ with a logistic growth function to capture the cumulative arrival of birds (Fig S 2). The population is assumed to be mixed homogeneously. The logistic cumulative arrival model captures realistic population size fluctuation over time during the breeding season by modelling arriving birds over time as a logistic growth function fitted with demographic data (Fig S 3). The sigmoid increase in alive birds reflects cumulative arrivals, distributed around a mean arrival date. As discussed below, the changing population size is key to modelling transmission dynamics of wildlife diseases and yet is often difficult to infer due to lack of time series demographic data^(9)^. The model is defined by Equations 1.

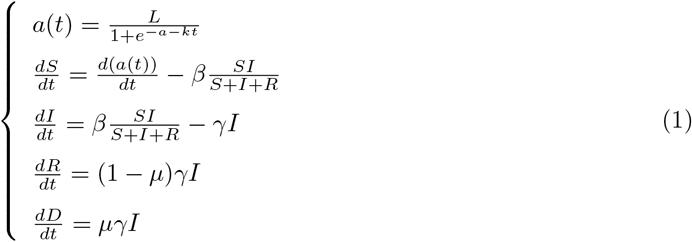

where *β* is transmissibility, 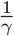 is infectious period and *μ* is mortality ratio of infectious individuals. These epidemiological parameters are estimated using maximum likelihood estimation, implemented in R package ‘bbmle’^(63)^. Errors are assumed to be negative binomially distributed^(64)^. To avoid finding local rather than global minima, initial values of parameters are varied and two different optimization algorithms are used to ensure robust optimal values, following previously suggested practices^(64)^.

*a*(*t*) is the logistic growth function modelling pelican arrivals; *L* is the carrying capacity, *k* represents the steepness of the cumulative arrival curve; finally, *m* defines the midpoint of the curve where the growth rate peaks. These parameters (Extended data table) are estimated by non-linear least squares^(65,66)^. The logistic growth function of 2022 is fitted to the sum of alive bird counts and death bird counts, whereas *a*(*t*) of 2020 and 2021 are fitted to nest counts, as the nest counts in 2022 cannot reflect the real population size due to the devastating mortality of HPAI infections (Fig S 3). While we assume all the arrivals after the initial time are susceptible, initial number of infectious *I*(*t*_0_) are assumed to be proportional to the population size of the initial time 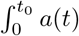, and therefore the initial number of susceptibles are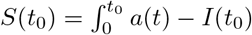.

#### Modelling vaccination

Vaccination is modelled in the simplest form^(61)^: assuming a perfect vaccine that prevents infection; the vaccinial immunity is assumed to last for the whole breeding season; the vaccine is introduced immediately before the outbreak start time. The initial infectious individual is assumed to be 1, and the vaccinated individuals are added to Recovered compartment in the model. The model is essentially the same as Equations 1; the only differences are the initial conditions of susceptible, infectious and recovered individuals, defined as Model 2, where *ρ* is ratio of vaccinated birds of the initial susceptibles (Fig S 2A).

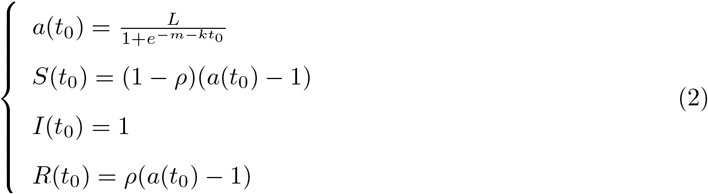

#### Modelling potential transmission from dead birds

To evaluate the impacts of carcass removal during 2022 outbreak, we extend the simple SIR model to include transmission from carcasses (Fig S 2B), defined by Equation 3. The model assumes that dead birds are infectious on average of 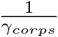 days, represented by *D*_*inf*_. Afterwards, they degrade and become non-infectious, represented by *D*_*deg*_.

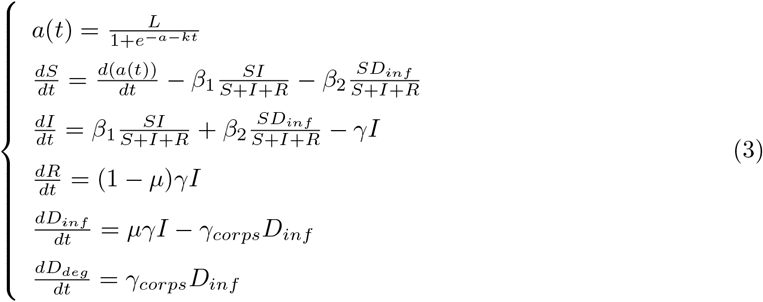

where *β*_1_ is transmissibility of infectious alive birds and *β*_2_ is transmissibility of infectious dead birds. The carcass removals are conducted respectively at Day 75, 76, 77, 79, 82, 100, 104 (Julian dates) of 2022. The simulation runs through the days until the carcasses are removed, and the number individuals are then removed from *D*_*inf*_ and *D*_*deg*_ compartments based on ratio of 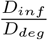 on the dates, reflecting the realistic removal process, and then, the simulation runs onward with the new initial conditions until Day 120.

## Data and code availability

All code and data are provided on https://github.com/kikiyang/h5-dp-gr.

## Acknowledgments

We acknowledge assistance of Christiana Mavridou, DVM in sample collection from Dalmatian pelicans. We acknowledge contribution of Haris Nikolaou in population monitoring of Dalmatian pelicans. We acknowledge field assistance of staff members at the Society for the Protection of Prespa, Hellenic Ornithological Society, and the Natural Environment and Climate Change Agency of Greece. We acknowledgment contribution of George Georgiades and other members of National Reference Laboratory for Avian Influenza and Newcastle Disease at Department of Avian, Honey Bee and Aquatic Organisms Diseases, Directorate of Thessaloniki Veterinary Center, Directorate General of Veterinary Services, Ministry of Rural Development and Food in Greece for providing sequencing results from outbreaks in Greece. S.M. acknowledges support by INVESTIGO grant (No. 1769562055/2022-INVGO-12) from the Spanish Ministry for Research, Innovation and Universities. B.T.G. was supported by the Princeton Catalysis Initiative and Princeton Precision Health. We acknowledge funding from the Society for the Protection of Prespa, through support provided by the Hans Wilsdorf Foundation. This material is based upon work supported by the High Meadows Environmental Institute at Princeton University through the Mary and Randall Hack ‘69 Research Fund.

## Author contributions

Q.Y., O.A., B.T.G., and G.C. conceived and planned the research. O.A. and G.C. curated population monitoring, movement tracking, and disease outbreak data. Q.Y., O.A., and G.C. collected samples in the field. U.H., S.M., S.C.C., E.M., C.I.D. conducted diagnoses of samples. B.T.G. advised on analysis methodologies. Q.Y. performed the analyses and wrote the initial manuscript. L.M. advised on interpretation of phylogenetic analyses. All authors edited and approved the manuscript.

## Supporting Information captions

Table S1: Fitted parameters for a logistic growth model of 2020, 2021 and 2022 arrivals.

Table S2: Summary of influenza virus detection and serology survey of captured Dalmatian pelicans.

Fig S1: Counts of background sequences collected in each country in the phylogenetic analyses.

Fig S2: The SIR epidemiological models for 2021 and 2022 outbreaks (A, without vaccination), with arrivals modelled as a logistic growth functions in Fig S 3, and the models for simulating vaccination (A) and carcass removal (B).

Fig S3: Logistic growth modelling of migratory arrivals in 2020, 2021 and 2022. The black curves are fitted number of arrived birds, with 95% confidence interval shown in pale blue area. The red curves are modelled arrival rates. The blue points are observed arrived birds, which are represented by nest counts in 2020 and 2021 and by the sum of dead and alive birds in 2022.

Fig S4: Movement trajectories of two Dalmatian pelicans (‘Adriatiku’ and ‘Albella’) that travelled between Albanian colony (Karavasta lagoon) and Prespa colony (Lesser Prespa Lake) (Movebank study ID: 4108676183) and one Dalmatian pelican (‘Golub’) that travelled between Skadar Lake colony in Montenergo, Albanian colony and Prespa colony (Movebank ID: 6750250558).

Fig S5: Dalmatian pelican death counts on Prespa colony during the 2021 and 2022 outbreaks.

Fig S6: Bayesian time-scaled phylogeny of 4351 HA sequences. The pelican symbols highlight sequences sampled from outbreaks in pelican species with mortality of more than 1,000. Besides the outbreak in Greece (Outbreak #3), Outbreak #2 is in Dalmatian pelicans in Russia and Outbreak #1 is in great white pelicans in Senegal and Mauritania. Coloured circles represent sequences sampled from pelican species.

Fig S7: Maximum likelihood trees of internal genes.

